# Bumblebees locate goals in 3D with absolute height estimation from ventral optic flow

**DOI:** 10.1101/2024.10.14.618154

**Authors:** Annkathrin Sonntag, Mathieu Lihoreau, Olivier Bertrand, Martin Egelhaaf

## Abstract

Bumblebees rely on visual memories acquired during the first outbound flights to relocate their nest. While these learning flights have been extensively studied in sparse environments with few objects, little is known about how bees adapt their flight in more dense, cluttered, settings that better mimic their natural habitats. Here we investigated how environmental complexity influences the first outbound flights of bumblebees. In a large arena we tracked the bees’ 3D positions to examine the flight patterns, body orientations, and nest fixations across environmental conditions characterised by different object constellations around the nest entrance. In cluttered environments, bees prioritised altitude gain over horizontal distance, suggesting a strategy to overcome obstacles and visual clutter. Body orientation patterns became more diverse in dense environments, indicating a balance between nest-oriented learning and obstacle avoidance. Notably, bees consistently preferred to fixate the location of the nest entrance from elevated positions above the dense environment across all conditions. Our results reveal significant changes in the 3D flight structure, body orientations, and nest fixation behaviours as object density increases. This highlights the importance of considering 3D space and environmental complexity in understanding insect navigation.

## 1 Introduction

Bees use visual memories to return to their nest after foraging trips. These visual memories are thought to be acquired during the first outbound flights when the bees do not forage but perform convoluted manoeuvres consisting of many loops and arcs (e.g. Capaldi et al. (2000), Philippides et al. (2013), and Woodgate et al. (2016); for reviews see: Collett et al. (2018) and Collett et al. (2023a)). During these first flights, the bees use a “turn back and look” behaviour to regularly face the nest and presumably learn the visual surroundings (Lehrer, 1991; Lehrer, 1993). Bees must perform these learning flights to efficiently return to their nest and avoid risky, time-consuming searches at the end of foraging bouts (Degen et al., 2016). After a short walking phase, the bees gradually increase their flight altitude and distance to the nest by progressively enlarging the size of their loops, thereby expanding the area covered by these flights (Capaldi et al., 2000; Osborne et al., 2013; Woodgate et al., 2016; Lobecke et al., 2018; Bertrand et al., 2023). Typically, the learning flights were investigated at a detailed level in simple environments with only few cylindrical objects surrounding the nest (Hempel de Ibarra et al., 2009; Collett et al., 2013; Robert et al., 2018; Collett et al., 2023b). For instance, it was found that bumblebees adjust their body orientation toward specific directions such as toward directions relative to the geomagnetic north or towards visual features in the surroundings (Hempel de Ibarra et al., 2009; Collett et al., 2013; Collett et al., 2023b). Additionally, studies have focused on the coordination between head and body movements during these flights (Odenthal et al., 2020; Doussot et al., 2021).

In nature, however, bee nests are often found in cluttered environments such as grasslands, forests, or agricultural cropland (O’Connor et al., 2017; Liczner et al., 2019). These natural habitats may present obstacles that bees must overcome or navigate around. Additionally, visual clutter and occlusions complicate orientation by eliminating reliable visual cues like a visual compass and making it harder to learn the environment. Consequently, dense and cluttered environments may require bees to develop more specific learning and navigation strategies. While the development of the flight altitude played a minor role in aforementioned studies in sparse environments, the described challenges in more ecologically realistic dense environments highlight the importance of understanding how the 3D structure of learning flights are affected.

Here, we investigated how the features of the environment shape the 3D characteristics of the first outbound flights of bumblebees in the immediate vicinity of their nest hole. Objects surrounding the nest entrance may serve as landmarks indicating the nest’s position, but they also present challenges such as occlusion, higher collision risks, and dramatic visual changes when transitioning from a dense environment to a more open one. Therefore, we examined how during the initial outbound flights the increase in altitude and distance from the nest is influenced by the features of the surroundings such as the density of these objects’ constellations, and the distance between the objects and the nest. In denser environments, the bees could increase their horizontal distance to the nest while keeping their flight altitude low (small altitude distance ratio) as we found in a previous study that homing bees prefer to enter dense environments at low altitudes (Sonntag et al., 2024). Alternatively, the bees could increase their flight altitude while keeping the distance to the nest small (large altitude distance ratio) because the bees may try to ‘escape’ the clutter to gain an overview of the environment. Such altitude gain in dense environments is expected by homing models based on visual memories (Sonntag et al., 2024). We also investigated the relationship between the positions where bees fixate on the nest entrance — possibly acquiring the visual memories necessary for their return—and their flight altitude in relation to the height of surrounding objects. In sparse environments bees were reported to show a clear body orientation towards the nest location while the flight direction deviated from this direction (Collett et al., 2013; Philippides et al., 2013; Collett et al., 2023b). A dense environment will pose a challenge in keeping the body axis oriented towards the nest due to a higher collision risk and might thus interrupt fixation behaviour. In addition, the body orientation and flight direction could change with altitude when they are affected by the visual clearance or occlusion of the objects.

## 2 Methods

### 2.1 Animal handling

We sequentially used three healthy hives of *Bombus terrestris* provided by Koppert B.V., The Netherlands. After arrival, the bees were transferred under red light (non-visible to bees (Skorupski et al., 2007)) into an acrylic box (30×30×30cm). This box was covered with black cloth to mimic the natural, underground nesting conditions of *B. terrestris* (Goulson, 2010). The nest box was connected to the flight arena via a system of six small boxes (8cmx8cmx8cm) and plastic tubes (2.5cm in diameter). One of the boxes contained a micro-gravity feeder that gave the hive direct access to sugar solution. The feeder consisted of a bottle with a small plate at the bottom where the bees could feed on sugar solution (30% sugar in volume) ad libitum through small slits in the plate. Pollen balls (50ml ground, commercial pollen collected by honeybees (W. Seib, Germany) and 10ml water) were provided ad libitum directly into the nest box. After their first outbound flight, the bees were marked with numbered plastic tags glued on their thorax with a melted resin. The temperature in the experimental room was constantly kept at 20° degrees and artificial light from above was provided in a 12h/12h day-night cycle.

### 2.2 Experimental Design

The flight arena was similar to Sonntag et al. (2024) (see Fig.1A). It consisted of a cylinder with 1.5m in diameter and 0.8m in height. The walls and the floor were covered with a red and white pattern (perceived as dark and white by the bees Skorupski et al. (2007)) with a spatial frequency distribution of 1/f characteristic of natural sceneries (Schwegmann et al., 2014). This pattern provided enough contrast for the bees to use optic flow for flight control but did not give any directional information for navigation. To facilitate the video recording and tracking of the bees, the arena was lit from below with 18 neon tubes (36W Osram + Triconic, light spectrum in Supplementary Material). This light was filtered by a red acrylic plate (Antiflex ac red 1600 ttv) so that the bees were undisturbed by the lighting outside their perceptual range (Skorupski et al., 2007). The arena was covered with a transparent acrylic ceiling at a height of 0.6m. The ceiling allowed lighting from above by 8 neon tubes (52 W Osram + Triconic) and 8 LEDs (5W GreenLED, light spectrum in Supplementary Material; as in Sonntag et al. (2024)), and recording via six high-speed cameras with different viewing angles.

**Figure 1.**
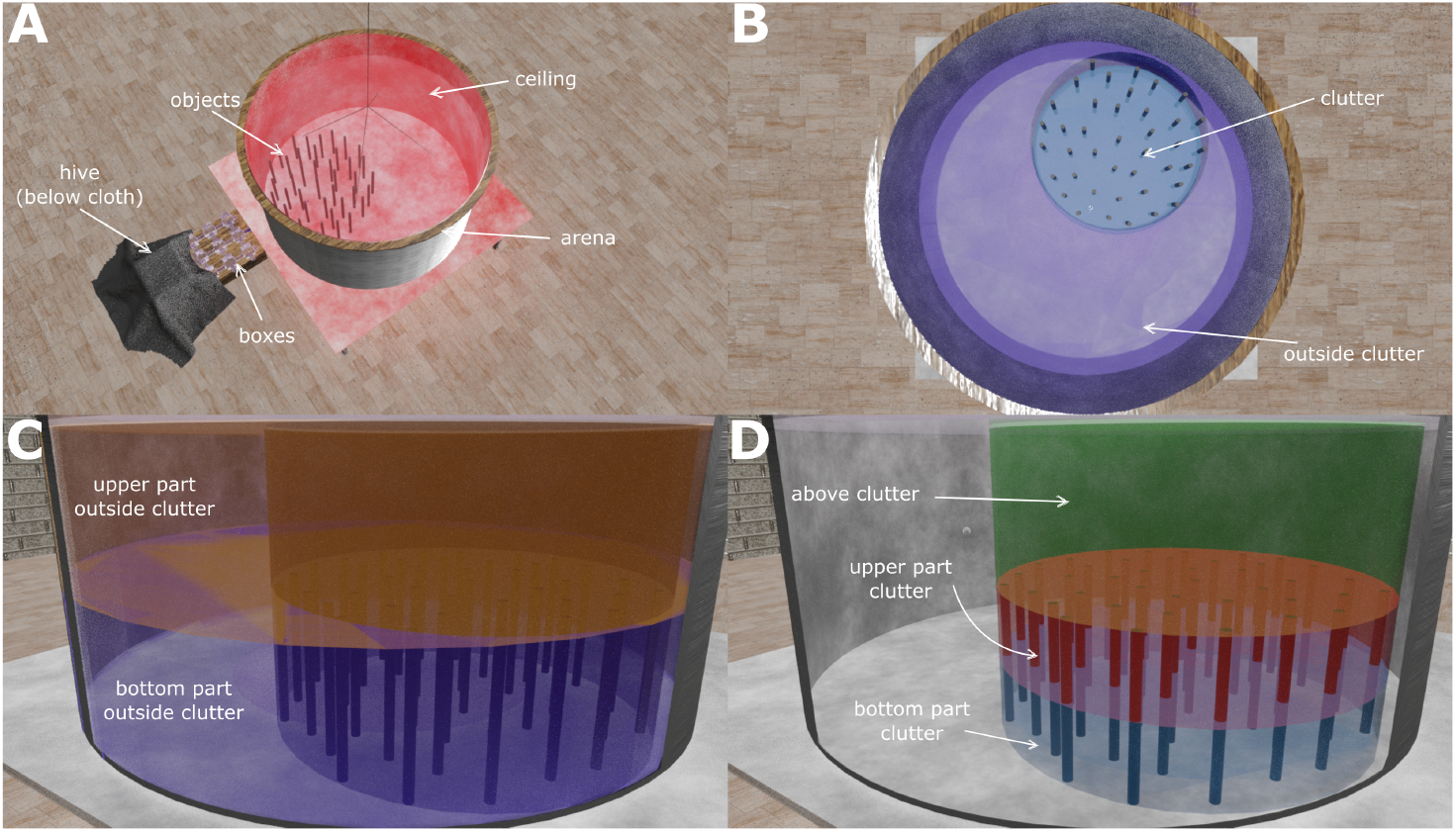
The setup with the flight arena and the spatial categories used to split the flight volumes for the comparison of the nest fixations. **A:** The hive of the bumblebees was connected to the cylindrical flight arena (1.5m in diameter, 0.6m in height) via a system of small boxes and tubes to select one bee at a time. A selected bee could enter the arena through a tube from below. The entrance hole was surrounded by objects (the number and arrangement depending on the tested environment). Six cameras recorded the bees’ position and orientation (not shown in the image). **B:** To compare the flight volumes where the bees fixated the nest, we split the arena in two main categories: the cluttered area around the nest (clutter, indicated by the blue circle) and the area outside the clutter (indicated by purple). **C:** The height outside the clutter was then split into the bottom part including low altitudes (BOC) and upper part including high altitude (UOC). **D:** The height of the cluttered area was split into three altitudes: the low altitude bottom part of the clutter (BC), intermediate altitude at the upper part of the clutter (UC) and above the clutter (AC).

A bee crossed two of the small boxes to enter the arena through a hole from below through a plastic tube (2.5cm in diameter). We used these boxes to separate single bees for the recordings while the other bees could forage undisturbed. The entrance to the flight arena was surrounded by red, cylindrical objects (30cm in height and 2cm in diameter), the exact number and positions depended on the test condition. We used four environments by varying the number and arrangement of objects in the arena: The 1st environment, “three objects environment” was sparse with three objects arranged close to the nest entrance (distance ¡ 0.1m, Fig. 2A) similar to previously tested environments for learning flights (Lobecke et al., 2018; Robert et al., 2018; Doussot et al., 2021). The three other environments varied in number and density of the objects and covered a circular area around the nest entrance with a diameter of 0.8m. The 2nd environment of our investigation, “full-density”, consisted of 40 randomly placed objects (Fig. 2B, the three nearest objects to the nest were the same as in the “three object” test) that provided a challenge for the bees but it provided enough space for the bees to fly through (Gonsek et al., 2021; Sonntag et al., 2024).

**Figure 2.**
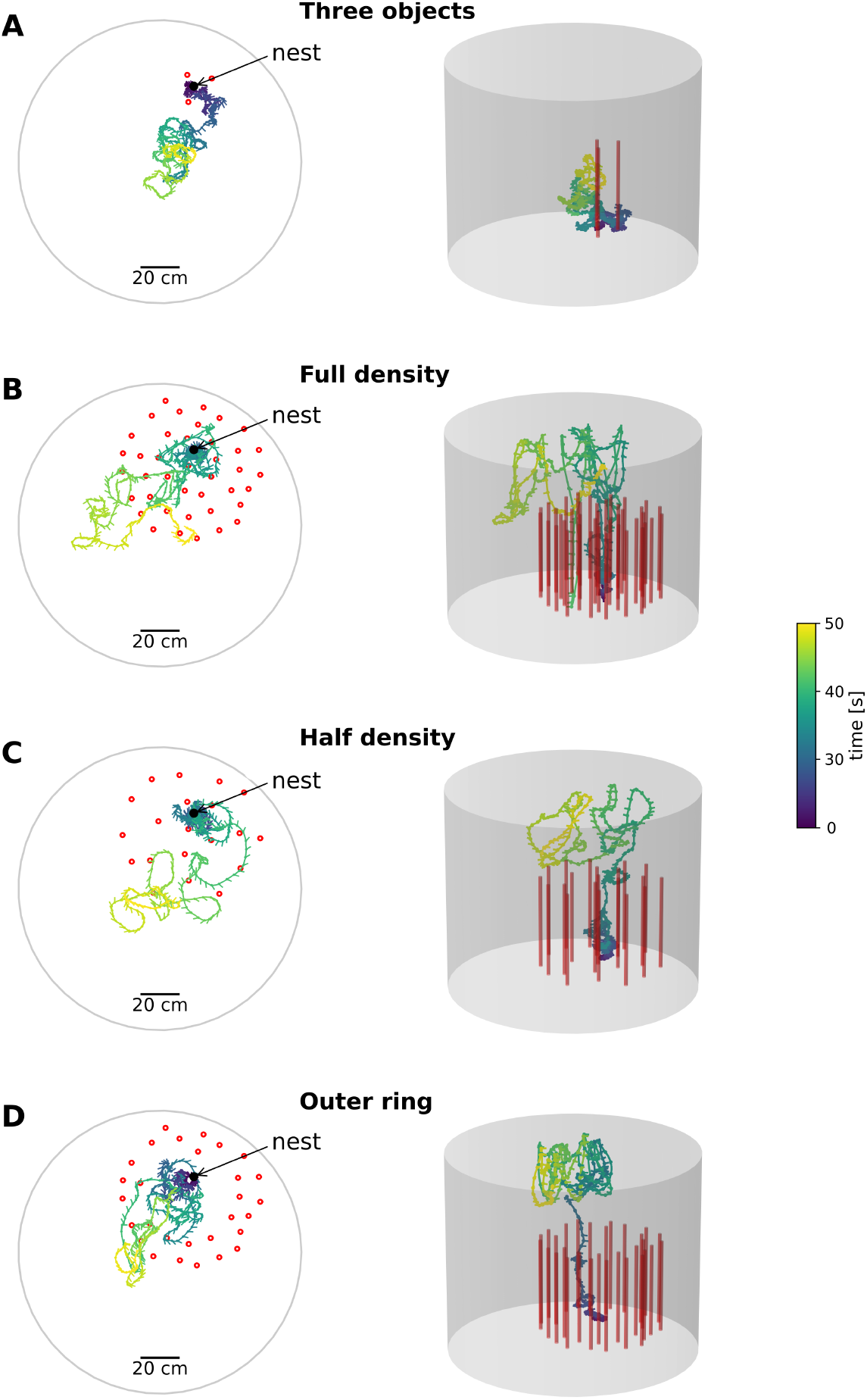
Exemplary flights in the four environments. The trajectories are colour coded by the time, blue indicating the entry to the area and yellow after 30 seconds of flight after take-off in addition to an initial walking period (leading to varying durations). The objects are indicated by red circles in the 2D plots (left column) and red cylinders in the 3D plots (right column). The nest indicated by an arrow is the nest entrance to the flight arena. **A:** Three objects environment: three objects surrounding the nest. **B:** full-density environment: 40 Objects surrounding the nest. **C:** half-density environment: 20 objects surrounding the nest. **D:** Outer ring environment: A ring of 30 objects surrounding the nest with the same density as with 40 objects.

To generate the placement of the objects we used another experiment as a reference where the behaviour of bees was recorded in a tunnel containing 49 10×3×300mm3 objects (Gonsek et al., 2021). In our study, we used cylindrical objects that have a base with a diameter of 22mm diameter. We randomly placed the objects incrementally, whereby the distance between neighboring objects had to be at least 9 cm; otherwise a new location had to be generated for the object. We generated multiple object constellations containing the 40 objects. Since we knew the bee navigated well in the experiment (Gonsek et al., 2021), we selected a distribution of objects closest to the distribution of objects in Gonsek et al. (2021). This was accomplished by choosing the constellation having the closest distribution of distance between neighbouring objects.

The 3rd environment, “half-density”, consisted of 20 objects (Fig. 2C, the three nearest objects to the nest were the same as in the “three objects” test) with half the density of the “full-density” environment.In this object arrangement, we removed 20 objects to create distance distributions between the four nearest objects (first closest, second closest, and so on), resulting in distances that are half the average in the full-density arrangement. The 4th environment, “outer ring”, consisted of 30 objects at the border of the circular area around the nest with the same density as in the “full-density” environment (Fig. 2D). While the “outer ring” environment tested an increased distance between the nest and the objects, it also yields similarities to Niko Tinbergen’s famous pine cone ring to test visual learning in wasps (Tinbergen, 1932; Tinbergen et al., 1938). Throughout the tests, the objects were initially arranged as in the “full-density” setup. For the “three objects,” “half-density,” and “outer ring” environments, a subset of this original object arrangement was used. In the “full-density” and “half-density” environments we challenged the bees with different densities but a similar area that was covered. In contrast, in the “outer ring” environment we tested if the density alone influenced the behaviour or if it is also influenced by the distance of the objects to the nest entrance. The cylindrical arena could be accessed through a door to modify the object arrangements for different environmental conditions. Each bee was tested in only one of the four environments. In total, we recorded 22 bees per environment, resulting in 88 unique individuals tested (colony distribution in Supplementary Material). Only one bee at a time was allowed to enter the flight arena. Between the flights, the arena was cleaned with 70% Ethanol to remove potential chemical markings (Cederberg, 1977; Foster et al., 1989; Chittka et al., 1999; Eckel et al., 2023). Each bee was recorded for 2min after entering the arena. If a bee took longer than 1min to take-off, the bee was captured, released back to the hive and the recording was discarded.

### 2.3 3D Flight trajectories

The bee trajectories in the arena were recorded at a frequency of 62.5 Hz (16 ms between consecutive frames) using six synchronised Basler cameras (Basler acA 2040um-NIR) positioned at different angles, similar to previous studies (Doussot et al., 2021; Eckel et al., 2023; Sonntag et al., 2024). One camera was mounted above the centre of the arena to capture the bees’ planar movements. Another one was positioned above the centre of the cluttered area around the nest, while the remaining four cameras were arranged around the arena to record the bees’ positions from different perspectives, reducing triangulation errors in 3D positioning. Recording commenced prior to the bees’ entry into the setup, and the initial 60 frames were utilised to generate a background image of the arena. Only image sections (40×40 pixels) with significant differences from the background image were saved. These sections likely indicated the presence of a bee and were stored along with their coordinates. The recording scripts were written in C++. A custom neural network is used to analyse image sections to determine whether they contain bees. The network was a five layer feed-forward network. The first three layers were convolutional with 40, 64, and 32 filters respectively. The fourth layer was a dense layer. Finally the last layer was single neuron encoding the probability of the input to be a bee or not a bee. The activation functions used were a linear rectifier (Re-Lu). We trained the model for 10 epochs and reached an accuracy of nearly 100% after training. We trained the model on roughly 1Mio manually classified images. The trained model along with the database and model pipeline (from labeling the images for training, training the model, evaluating it and applying the model to classify images) is accessible in a repository and data publication (Bertrand et al., 2024b; Bertrand et al., 2024a). Crops with non-biological speeds (above 4 m/s Goulson (2010)) or implausible positions (outside the arena) were manually reviewed. The trajectories were analysed using Python (version 3.8.17), primarily with OpenCV with a custom written analysis pipeline (Bertrand et al. (2024c), version 0.2.0). DeepLabCut, a software widely used for markerless pose estimation in animal studies, was used to identify the head and abdomen positions to determine the orientation of the bee’s body-length axis (Nath et al., 2019). A comprehensive list of the packages used is available in a data publication (Sonntag, 2024).

### 2.4 Statistical analysis for hypothesis testing

To investigate how the environment influenced the structure of the learning flights of bees, we calculated the ratio between the bee’s altitude and the 2D distance of the bee to the nest (x-y plane). We hypothesised that in denser environments, bees might increase their horizontal distance from the nest while keeping a low flight altitude, mimicking their homing behaviour (Sonntag et al., 2024). Alternatively, bees in denser environments might increase their flight altitude while staying close to the nest. This could help them gain a better overview of their surroundings, as suggested by homing models based on visual memories (Sonntag et al., 2024). We calculated the mean ratio for 10s after the first take-off (at least 0.01m above the ground) and excluded positions where the bees were walking (z ¡0.01m). Before the calculation of the mean ratio, we smoothed the altitude and horizontal distance by calculating the rolling mean over a time window of 250ms. We focused on the initial altitude increase before bees began descending or hitting the ceiling, choosing a 10-second window when most bees showed their typical altitude increase (see Supplementary Material). We also validated that the results were independent of the time window by calculating the altitude-distance ratio for windows between 5 and 30 seconds. For a statistical comparison of the altitude distance ratios we used the Kruskal Wallis test and the post-hoc Dunn test because the data was not normally distributed. In addition, we used a linear regression to investigate the effect of increasing object number on the altitude distance ratio. We coded the conditions (three objects, half-density with 20 objects, outer-ring with 30 objects and full density with 40 objects) as an ordinal variable from 0 to 3 because we are interested in the qualitative changes induced by the spatial arrangements rather than the absolute number of objects.

We hypothesised that the orientation of the bees’ body and their flight direction would be influenced by the object constellations near the nest. Furthermore, the body orientation and flight direction could be adapted differently at different altitudes if they were influenced by the visual clearance or occlusion of the objects. Therefore, we looked at the yaw angle of the bees relative to the nest during the first 30 seconds after the first take-off (at least 0.01m above the ground) separated into five categories: bottom part of the clutter (BC), z ¡ 0.15m; upper part of the clutter (UC), 0.15m =¡ z ¡ 0.3m; high altitude above clutter (AC), z ¿= 0.3; bottom part outside the clutter (BOC), z ¡ 0.3m; upper level outside the clutter (UOC), z ¿= 0.3m (Fig. 1B-D). We used the area of the clutter for all conditions to compare similar regions between all conditions (Fig. 1B). For visualisation of the body orientation and flight direction we used circular kernel density estimations. The bandwidth of the circular kernels, which controls the degree of smoothing applied to the data, was estimated as 191 using a Python adaptation of the bandwidth estimation function from the R package “circular”. Further, the Rayleigh test was used to check whether the bees’ body orientations were uniformly distributed, while the v-test assessed whether they were oriented toward the nest, a specified mean direction (Landler et al., 2018). To test if the distributions of the body orientation are dependent on the environmental condition and the spatial categories, we used the two-sample Kuiper test.

Then we calculated the fixations of the nest for the full duration of the recorded learning flight (we only excluded positions 5cm below the ceiling to remove possible positions where the bees collided with the ceiling) similar to Robert et al. (2018) (Fig. 6). The nest fixations are defined as the times the nest position was stationary on the retina (Robert et al., 2018). Frames with a yaw angle of *±* 12.5 deg relative to the nest entrance were taken as one sample. The following frames were added to this sample if the absolute angle difference was smaller than 2.4 deg (150 deg/second as in (Robert et al., 2018)). One sample was only valid if it contained at least 5 consecutive frames (80ms, as in Robert et al. (2018))). These fixations were separated into the same five altitude sections (BC, UC, AC, BOC, UOC). We used a two-way ANOVA to compare nest fixations across altitude sections, both within individual conditions and between different conditions. In addition, we used the Tukey HSD post-hoc test for pairwise comparisons of each altitude section. However, because the two-way ANOVA revealed a significant interaction between altitude sections and conditions, interpreting the main effects alone may be misleading. To address this, we performed an estimated marginal means (EMM) analysis, which provides the predicted means adjusted for the interaction effects. This method allows for a more precise comparison of nest fixation frequencies by accounting for the variability introduced by the interaction between factors, ensuring robust and interpretable results. The confidence intervals for the EMMs were also computed to provide insight into the reliability of these estimates. In addition we calculated the elevation angle between the bees body position and the nest position in the arena to check for the feasibility of the nest fixations. For all of these analyses python 3.8 and the packages statsmodels, bioinfokit and scipy were used.

## 3 Results

### 3.1 Bees increase flight altitude in dense environments

During learning flights, bees increase their altitude and distance to the nest (Capaldi et al., 2000; Osborne et al., 2013; Woodgate et al., 2016; Lobecke et al., 2018). We investigated how these flight characteristics are influenced by different constellations of objects. The initial part of the first learning flight was compared between the four environmental conditions (Fig. 3). When the nest entrance was surrounded by only three objects, an environment similar to other studies (Cartwright et al., 1983; Linander et al., 2018; Lobecke et al., 2018; Robert et al., 2018), we found a median altitude distance ratio of 0.61 (SD = 0.35) indicating that the bees increased more their 2D distance to the nest entrance than their altitude. In contrast, in the full-density environment we found a median altitude distance ratio of 1.73, indicating a larger increase in altitude than in 2D distance, and a larger variance (SD = 1.2). To determine whether the density of the objects, and thus the distance between them reflecting the degree of challenge of the environment for flight control, was responsible for this effect, we reduced the object density to half of the number of objects in the same area. In the half-density environment, we found a median altitude distance ratio of 1.15 indicating an increase of ratio of altitude to distance between the full-density environment and the environment with three objects. The variance with the half-density environment (SD = 0.8) was also in between that obtained for the full-density environment and the three object environment. Lastly, we tested how the distance of the objects to the nest influenced the altitude and distance increase. The outer ring environment resulted in intermediate values for both the median ratio (M = 1.16) and the variance (SD = 0.51) falling between those of the three-object and full-density environments.

**Figure 3.**
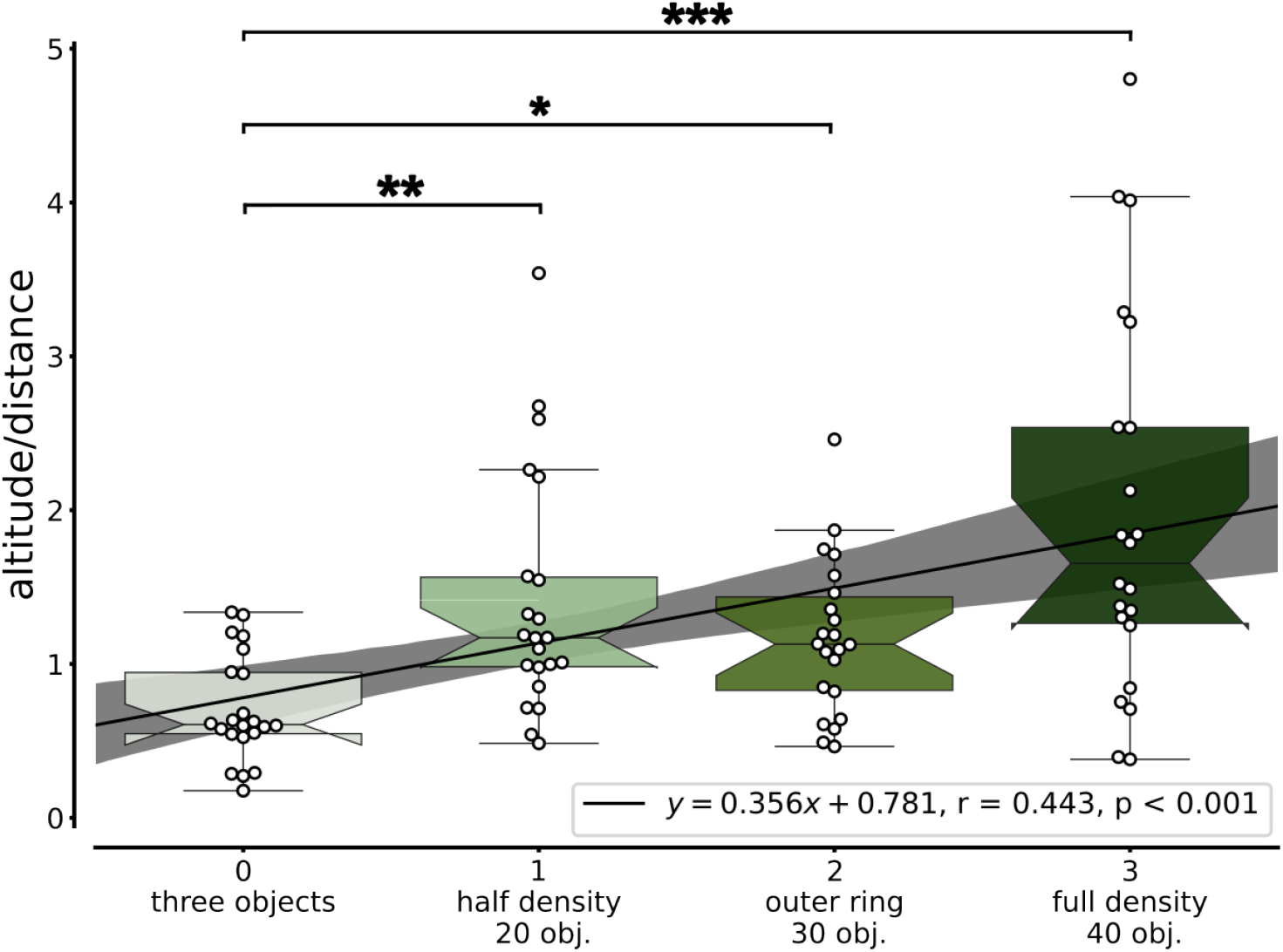
Altitude and distance ratio. Altitude and distance ratio for the initial part of the learning flights for the four environmental conditions: three objects, half-density (20 objects), outer ring (30 objects) and full-density (40 objects). The altitude distance ratio for each bee is shown as white circles. The hatched boxplots display the median and the whiskers show the lower and upper range of 1.5 times the interquartiles. The star code shows different levels of significance following the Dunn post hoc test. The ratios increased with increasing number of objects and differed statistically between the three object environment and the other environments (Kruskal Wallis test: H = 24.277, p *<* 0.001; Dunn’s post hoc test: p_full-density-three objects_ *<* 0.001, p_half-density - three objects_ = 0.006, p_outer ring - three objects_ = 0.043). The black line indicates the linear regression showing a increase of the altitude distance ration with increasing number of the objects. Ordinal condition numbers from 0 to 3 correspond to the conditions three objects, half-density (20 objects), outer-ring(30 objects) and full-density (40 objects). N = 22 bees per environmental condition.

A linear regression model could predict from the ordinal conditions the altitude distance ratio which means that the altitude distance ratio increases with increasing object number (p *<* 0.001, R = 0.443). Additionally, the altitude distance ratio differed between the “three objects” environment and all others (Kruskal Wallis test: H = 24.277, p ¡ 0.001; Dunn’s post hoc test: p_full-density-three objects_ *<* 0.001, p_half-density - three objects_ = 0.006, p_outer ring - three objects_ = 0.043). The other ratios were not statistically different from each other (Dunn’s post hoc test: p ¿ 0.05, Fig. 4). These findings are consistent for time windows between 10 and 30 seconds 4). These results demonstrate that the flight structure of the bees’ first learning flight depends on the environmental features around the nest, with bees increasing their altitude as the number of surrounding objects increases.

**Figure 4.**
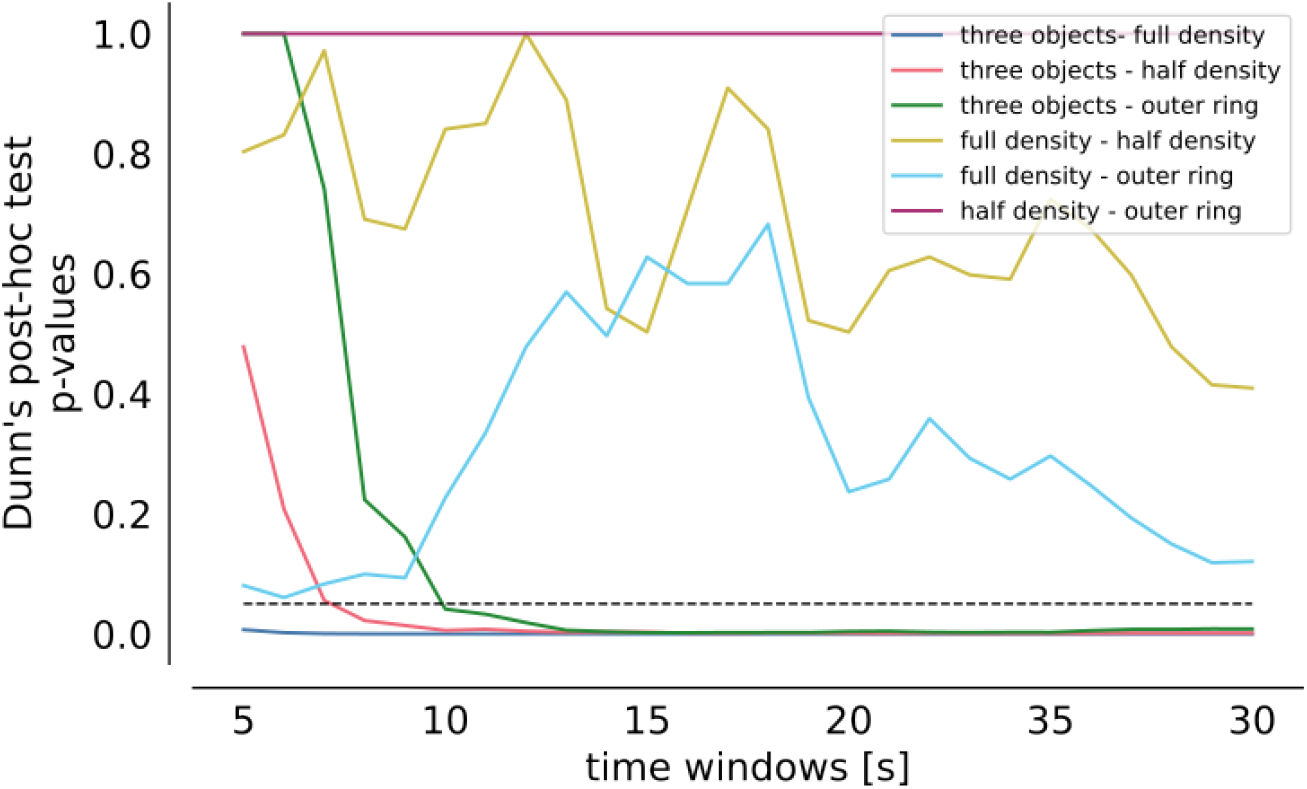
Dunn’s post-hoc test of the altitude distance ratio for varying time windows. Dunn’s post-hoc test p-values were calculated for varying time windows (5 to 30 seconds) to analyze the altitude and distance ratios during the initial part of the learning flights across the four environmental conditions: full-density, half-density, outer ring, and three objects. For time windows longer than 5 seconds, the ratios in the three-objects environment were consistently different from those in the other three environments. In contrast, the ratios for the other three environments remained similar across all tested time windows.

### 3.2 Bees orient towards the nest across all types of environments

During their first outbound flights, bees regularly turned back and looked toward the nest entrance (Lehrer, 1991; Lehrer, 1993). This behaviour results in deviations of the flight direction of the bees and the body orientation. Therefore, we analysed the yaw angle of the bee’s body relative to the nest entrance. The orientations were separated into five categories in regard to their altitude and 2D distance to the nest.

Statistical analysis revealed that all distributions of body orientation were non-uniform, except the distributions in the bottom part of the arena outside the clutter (BOC) in the half-density environment (Rayleigh test, see Supplementary Material). In all environments, the bees predominantly oriented toward the nest when flying within the cluttered area, at all flight altitudes (Fig. 5i-iii). The v-test revealed for these areas a mean body orientation of 0 degrees relative to the nest entrance (for details see Supplementary Material). Outside the cluttered area, the bees oriented mostly away from the nest, potentially to explore their surroundings (Fig. 5iv and v). Only in the half-density environment, the bees orient towards the nest when flying outside the cluttered area at a high altitude (v-test, p = 0.007).

**Figure 5.**
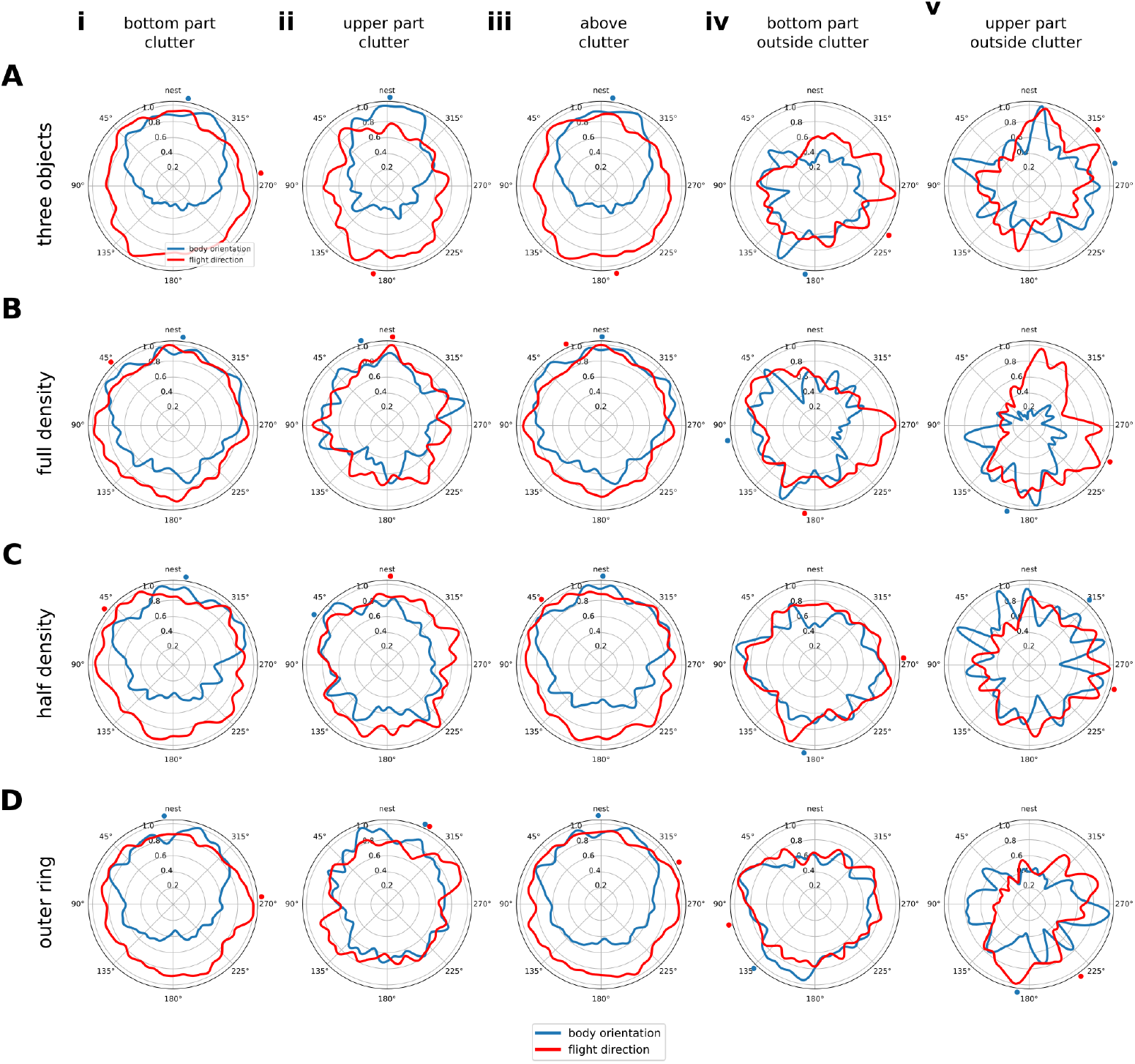
Body orientation and flight direction. Circular kernel density estimation distributions of the body orientation (blue line) of the bees along the yaw angle and their flight direction (red line) relative to the nest (0 deg) for the four environments (A, i: N (number of bees) = 22, n (number of data points) = 18536; ii: N = 22, n = 8236; iii: N = 22, n = 26772; iv N = 16, n = 3567; v: N = 13, n = 2360), full-density (B, i: N = 22, n = 13742; ii: N = 22, n = 4724; iii: N = 22, n = 18466; iv: N = 14, n = 4806, v: N = 9, n = 1970), half-density (C, i: N = 22, n = 11434; ii: N = 22, n = 4703; iii: N = 22; iv: n = 16137; N = 20; n = 8336; v: N = 16, n = 1830) and the outer ring (D, i: N = 22, n = 13606; ii: N = 22, n = 4963, iii: N = 22, n = 18569; iv: N = 21, n = 7393; v: N = 12, n = 2537). For each environment the body orientation and flight direction are shown for different layers in the area of the objects’ arrangement (clutter) (i bottom part clutter, ii upper part clutter and iii above the clutter) and outside the clutter (iv bottom part outside clutter, v upper part outside clutter). The mean direction is indicated by a dot at the outer edge of the polar plot for the body orientation (blue) and the flight direction (red). The star code gives the results of the v-test if the distribution has a mean direction towards the nest (0 deg).

The Kuiper test revealed that most body orientation distributions are different from the others (for details see Supplementary Material). However, in the full-density condition, the distributions of body orientations are similar within the area above the clutter compared to, respectively, low and intermediate within the clutter. In the three objects condition, the distributions within the areas low altitude clutter and high altitude outside clutter are similar to each other. Additionally, the body orientation distributions are similar when the bees fly at high altitudes outside the clutter in the three objects and half-density environments. The distributions in the outer ring and half-density environments are also similar within the high altitude arena area. The distributions are similar in the half-density environment at high altitudes outside the clutter and in the outer ring environment in the high altitude cluttered area. Lastly, the distributions are similar in the outer ring environment in the areas low altitude and above clutter.

In summary, our results demonstrate that bees predominantly oriented toward the nest entrance during their learning flights in all environments within the cluttered area, thus in close proximity to the nest location.

### 3.3 Bees fixate the nest more often when flying above the objects

To examine how the positions where the bees fixated the nest were influenced by their environment, we separated the bees’ nest entrance fixations (for a definition, see Methods) into spatial categories, as we did for the body orientations (6 and for more details see Supplementary Material). In the three-object environment, similar numbers of nest fixations occurred within and above the clutter (Fig. 6A). In the area outside the clutter, fewer nest fixations occurred. In the full-density environment, the bees primarily fixated on the nest while flying above the clutter. This was followed by the positions at the bottom part of the clutter and at the upper part outside the clutter. The fewest nest fixations occurred at the upper part outside the clutter. In the half-density and outer-ring environments, the nest fixations were distributed similarly to those in the full-density environment. Most nest fixations occurred within and above the clutter, primarily either above the clutter or at the bottom part. Bees oriented toward the nest with similar frequency both at the upper part of the clutter and outside the clutter at both altitude levels.

**Figure 6.**
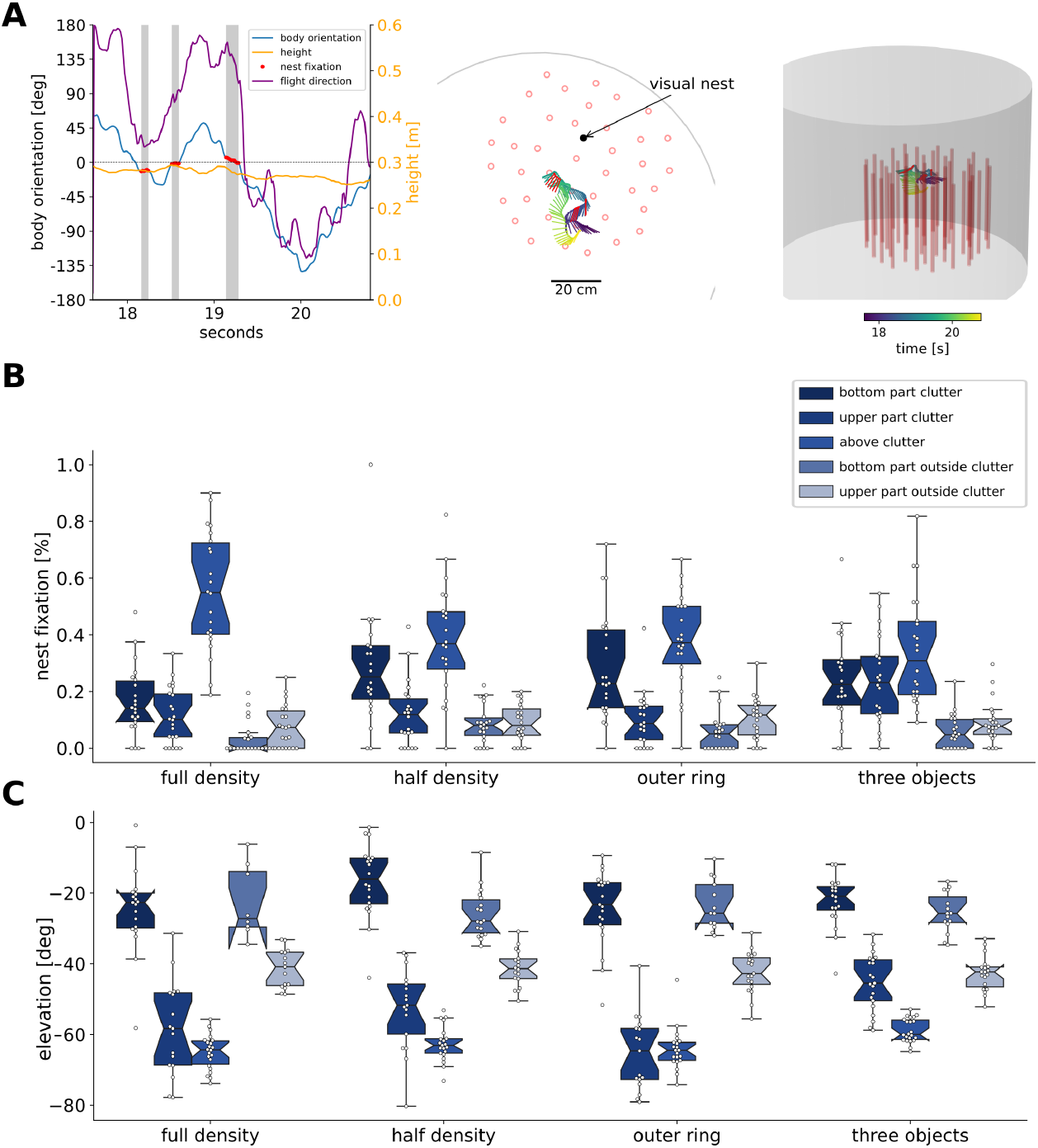
Nest fixations and their elevation angle. Example of how nest fixation were quantified (**A**). The left panel shows the times series of the body orientation (blue line), flight direction (purple line) and height (yellow line). The nest fixation block is indicated in the grey shaded area and red dots. The other two panels show the 2D (middle) and 3D (left) view of the exemplary part of the trajectory color coded by time. Nest fixations (**B**) and the elevation angle of these fixations (**C**) for five spatial areas (BC, UC, AC, BOC, UOC) in the four environments (full-density, outer-ring, half-density and three objects). Individual data are shown as white circles, hatched boxplots display the median and the whiskers show the lower and upper range of 1.5 times the interquartiles. N = 22 bees per environment.

Overall, we found a significant difference in nest fixation frequencies across the spatial categories (two-factor ANOVA, F(4) = 119.2, p ¡ 0.001), but no significant differences between the tested environments (two-factor ANOVA, F(3) = 0.165, p = 0.932). The interaction of the spatial categories and the environments was significant (F(12) = 5.098, p ¡ 0.001). We found that the bees fixated on the nest more often above the clutter than within the clutter or outside it (Tukey HSD post hoc test, p ¡ 0.001 for all comparisons). We also found more fixations within or above the clutter than outside it (Tukey HSD post hoc test, BC - BOC p ¡ 0.001, AC - UOC ¡ 0.001, AC - BOC = 0.007). However, we observed a trend suggesting that bees fixated on the nest more frequently when flying at within the upper part of the clutter compared to when flying outside the clutter at high altitudes, (Tukey HSD post hoc test, p = 0.088). Additionally, the nest fixations did not differ statistically between the bottom or upper part outside the clutter (Tukey HSD post hoc test, p = 0.584). The environmental conditions did not significantly influence the fixation proportions in the different spatial areas, but we found an interaction between the spatial categories and the environmental conditions (p ¿ 0.05 for all comparisons). The EMM of the nest fixations show that the interaction is mostly driven by the high proportion of fixation for the full-density environment in the area above the clutter while in the other conditions (especially the three objects condition) the proportion of fixation at the bottom part of the clutter is similarly high as in the area above the clutter (Fig. 7).

**Figure 7.**
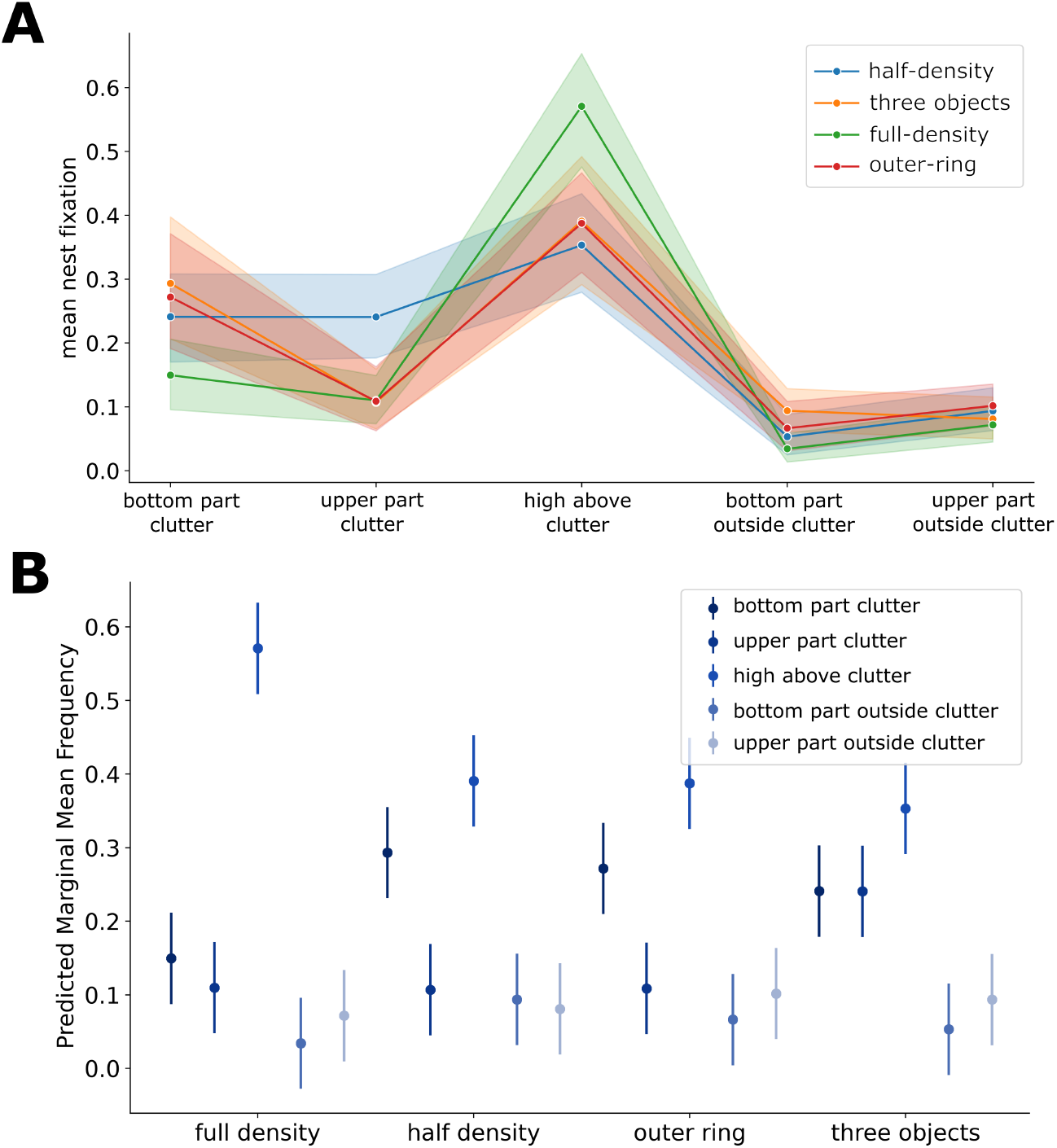
Interactions ANOVA and estimated marginal means. **A**) Interaction plot of the ANOVA of the mean nest fixation in the five spatial categories with the confidence interval shown with the shaded area around the line. (**B**) The estimated marginal means (with the 95% confidence interval) show that in the full-density environment most proportion of fixations occur in the area above the clutter while in all other environments the proportions are similar in the areas above clutter and low altitude clutter.

Since we could not determine the position and orientation of the bees’ head, we used their position relative to the nest in the arena as a proxy for the elevation angle of the nest within their visual field. An angle of zero degrees means that the body axis of the bees was horizontally aligned with the height of the nest, thus directly facing the nest. Decreasing elevation angles mean that the nest appeared lower within the bee’s field of view. Most of the elevation angles of the nest within the field of view of the bees ranged from 0 to -60 degrees while some outliers are found ¡ -60 degrees, especially for intermediate and high flight altitude levels (Fig. 6). Overall, the bees fixated on the nest primarily at higher altitudes above the object constellations, with the fixation pattern influenced by the combination of spatial areas and environmental features.

## 4 Discussion

Insect learning flights have mostly been analyzed in sparse environments so far (Hempel de Ibarra et al., 2009; Collett et al., 2013; Philippides et al., 2013; Robert et al., 2018; Doussot et al., 2021; Collett et al., 2023b), which do not accurately represent the typical habitats around bumblebee nests (O’Connor et al., 2017; Liczner et al., 2019). By manipulating the object constellations around a nest entrance, we found that bees adjust their flight patterns in response to environmental challenges, prioritizing altitude gain in cluttered environments and modifying their orientation patterns to balance nest-directed learning with obstacle avoidance. We observed a shift toward greater altitude gain in cluttered environments and more diverse orientation patterns in dense environments compared to sparse ones. Bees consistently preferred to fixate on the nest from elevated positions above the object arrangements.

### 4.1 Bees increase flight altitude in dense environments

Our study revealed significant differences in how bees adjust their altitude and distance from the nest based on environmental complexity. In a sparse environment with only three objects, bees increased their horizontal distance from the nest slightly more than their altitude. Similarly, the environment in which Stürzl et al. (2016) analyzed the learning flights of wasps report a similar increase in altitude and distance, which aligns with our findings in sparse environments. However, in dense environments with more objects, the bees prioritized gaining altitude over increasing their 2D distance. This demonstrates that bees adapt their flight strategies in response to environmental constraints. In cluttered environments, gaining altitude may provide better vantage points for visual learning from above. Indeed, high-altitude views offer a broader perspective of the landscape, which bees have been shown to use for identifying ground-level landmarks and navigating over long distances (Collett et al., 2015; Degen et al., 2016; Menzel et al., 2019; Brebner et al., 2021). Flying above the clutter may also reduce the need for obstacle avoidance. The increase in flight altitude observed in our study might therefore also be influenced by changes in ventral optic flow, which varies with environmental features. Previous studies have shown that bees adjust their altitude in response to ventral spatial textures (Linander et al., 2018). The higher variance in altitude-distance ratios in complex environments, compared to the sparse environment, may suggest that bees adopt more diverse strategies when navigating challenging landscapes. The similar altitude distance ratios in the half-density and outer ring environments need further investigations with changing specifically the distance of the objects to the nest and the density of the object constellation.

### 4.2 Bees orient towards the nest across environments

We observed a nest-oriented body orientation pattern near the nest across different environmental conditions, consistent with findings from previous studies (Hempel de Ibarra et al., 2009; Collett et al., 2013; Philippides et al., 2013; Doussot et al., 2021; Collett et al., 2023b). This supports the hypothesis that bees rely on local visual cues to establish spatial memories, particularly during their initial outbound flights (Zeil, 2022). Notably, the consistency of nest orientation across all altitudes in these cluttered environments suggests that bees effectively integrate three-dimensional spatial information to exhibit nest-oriented behavior.

In contrast to these similarities, we identified significant differences in the distributions of body orientation across environmental conditions and altitudes. While few comparisons revealed similar distributions, one notable finding was that bees exhibited similar orientation patterns above cluttered areas across low and high altitudes in the full-density environment. In addition, the orientation distributions at high altitudes outside the clutter were similar between the three-object and half-density environments, suggesting that bees may employ analogous strategies when facing intermediate levels of visual complexity. Furthermore, the resemblance between the orientation pattern at high altitudes outside the clutter in the half-density environment and at high altitudes within the clutter in the outer-ring environment indicates that bees adjust their orientation behavior based on the spatial arrangement of visual cues, rather than solely the density of landmarks

### 4.3 Bees fixate the nest more often when flying above the objects

Across all our environmental conditions, bees showed a preference for fixating on the nest entrance when flying above the clutter, even though the nest entrance was hidden and its location may have been occluded by the clutter. This finding is particularly interesting as it suggests that bees use elevated positions to gain clearer views of the nest and its surroundings, potentially creating more reliable visual memories.

Fixations at high altitudes raise the question of where the nest position appeared in the bee’s field of view. A modeling study on honeybee vision indicated that the resolution of a bee’s eye decreases when observing wide angles, especially as the vertical angles between ommatidia become smaller (Stürzl et al., 2010). Additionally, a morphological study on bumblebees showed limitations in perception in the ventral direction (Taylor et al., 2019). Therefore, observations of nest fixations at very low elevation angles should be interpreted with caution.

Within the dense environment and the areas outside it, the elevation angles fell within the perceptible range at flight altitudes. However, for intermediate altitudes within the dense environment and above the objects, nest fixations should be interpreted cautiously, as they are near the perceivable edge. Some high-altitude fixations might be at the border of the bee’s visual field (Taylor et al., 2019) and require further evaluation. Since we only investigated the yaw angle of the body orientation of the bees, further studies including 3D tracking of the head position and orientation are needed (e.g., (Doussot et al., 2021; Hateren et al., 1999)).

In the dense environments we tested, occlusions of the nest position by objects were likely. It is therefore noteworthy that we still found nest fixations. The consistent pattern of more fixations within or above the cluttered area, compared to outside it, implies that bees focus their learning efforts in the immediate vicinity of the nest, even in complex cluttered environments.

### 4.4 Implications and future directions

Our findings demonstrate that bumblebees exhibit remarkable flexibility in their navigation strategies. They alter their flight patterns and orientations based on environmental features, while the fixation of the nest entrance depends on the combination of altitude and object density. The preference for nest fixations from elevated positions, especially in cluttered environments, suggests that bees adjust their flight altitude to acquire visual memories for their return flights, depending on environmental conditions. This adds new insights to our understanding of insect navigation, which has long focused on movements in two dimensions and in sparse environments (Buehlmann et al., 2020; Zeil, 2022).

The varied orientation patterns observed in cluttered environments indicate a dynamic balance between exploring new areas and maintaining familiarity with the nest location. This balance may be crucial for efficient foraging in complex natural habitats with dense clutter.

In a previous study (Sonntag et al., 2024), which examined the homing abilities of bumblebees in a dense environment, we found that snapshot models of local homing (Dittmar et al., 2010; Doussot et al., 2021; Zeil, 2022) suggest that snapshot views taken at high altitudes—above the objects surrounding the nest entrance—yield better performance than those taken at lower altitudes. Our analyses of the learning flights in the present study align with this modeling result, with an observed increase in altitude over distance in dense environments, as well as nest fixations at heights above the objects.

Interestingly, behavioral experiments on return flights to the nest entrance have shown that bees do not require views from high altitudes to successfully return to their nest (Sonntag et al., 2024). One possible explanation is that low-altitude flights are more direct and efficient for returning to the nest location. Bees may decrease their flight altitude during consecutive learning flights to study low-altitude views around the nest, gradually updating their memories. While our study focused on the first learning flight of naïve bees, future studies on consecutive learning flights in dense environments are needed to better understand how learning develops in complex surroundings. Investigating the energetic costs of different flight strategies in various environments could also provide insights into the efficiency of these adaptive behaviors. Our analysis of bumblebee learning flights underscores the importance of studying the navigational behaviors of flying insects in three dimensions and in more diverse settings.

## Conflict of Interest Statement

The authors declare that the research was conducted in the absence of any commercial or financial relationships that could be construed as a potential conflict of interest.

## Funding

We acknowledge the financial support of the 3DNaviBee project by the collaborative funding of the German Research Foundation (DFG) and the French National Research Agency (ANR) (reference code: EG 82/22-1). While writing, ML received support from the European Commission (ERC Consolidator - GA101002644).

## Acknowledgments

We would like to thank Yigit Yargili and Melissa Vera Finke for their help during the data collection.

## Data Availability Statement

The data-sets and analysis pipeline for this study can be found in the repository “BumblebeeLearn-ingFlights3D” (Sonntag, 2024)..

## Notes

https://gitlab.ub.uni-bielefeld.de/a.sonntag/bumblebeelearningflights3d

### Competing Interest Statement

The authors have declared no competing interest.

### Summary of Updates

We analysed the influence of the time window on the altitude distance ratio and found the same statistical results for time windows between 10 to 30 seconds. Figure 5 was revised as we changed from linear KDE to circular KDEs. Exemplary time series plots are provided and fixation events are depicted. (Figures 6A and S2). We added a figure showing the spatial categories in the arena (new Figure 1) and also relabelled the different categories to improve readability.

