## Supplementary Material for "Bumblebees locate goals in 3D with absolute height estimation from ventral optic flow"

#### 1 Supplementary figures

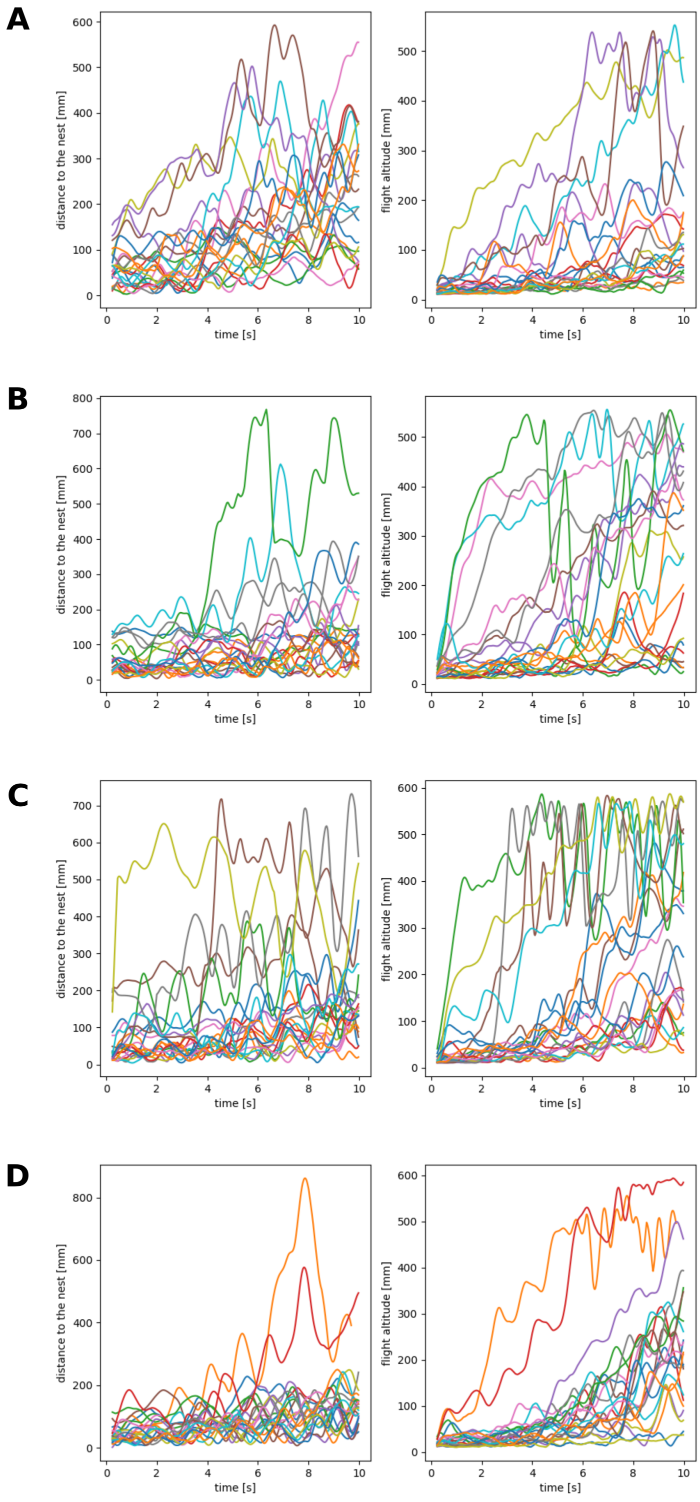


**Fig. S1**: **Distance to the nest and flight altitude.** Distance to the nest and height for each learning flight ($z$   0.01m) over time within the time window of 10 seconds after take-off for the four test conditions (A: three objects, B: full-density, C: half-density, D: outer ring).

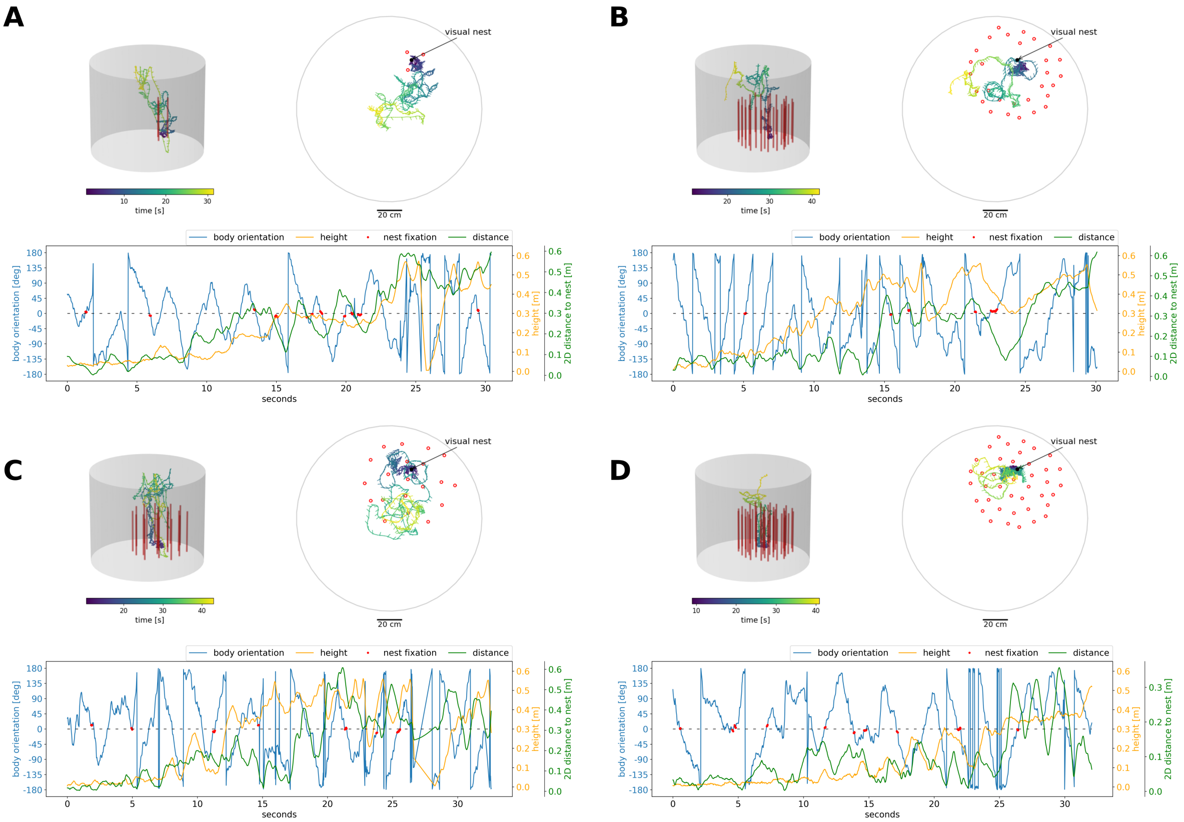


**Fig. S2**: **Exemplary trajectories.** Exemplary trajectories in 3D and 2D (top view) in the four tested environments (A, three objects, B: half-density, C: outer ring and D: full-density) with the body orientation, height and 2D distance to the nest for the initial walking phase and the first 30 seconds of flight. The fixations of the nest are highlighted in red.

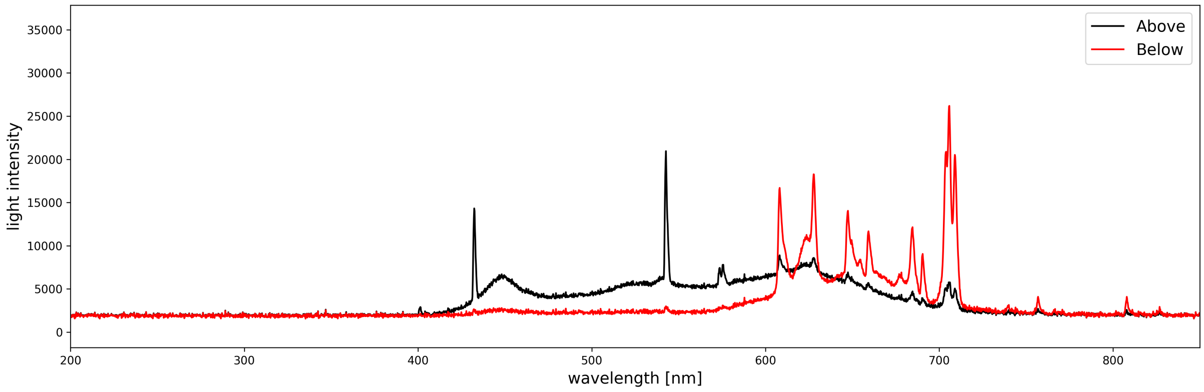


**Fig. S3**: **Light spectrum within the flight arena.** The light spectrum recorded within the flight arena from above (black) and from below (red). The light from above the arena ranges between 400 and 700 nm which is within the range of visible light to humans. The light from below, used for tracking the bee position, ranges between 600 and 700nm which is outside the perception of bees Skorupski, Döring, and Chittka (2007).

### 2 Supplementary tables

Table 1: Number of bees from colonies 1-3 in the four tested environments.

| hive | three objects | outer ring | half-density | full-density |
|---|---|---|---|---|
| 1 | 17 | 0 | 0 | 19 |
| 2 | 5 | 12 | 12 | 3 |
| 3 | 0 | 10 | 10 | 0 |
| total | 22 | 22 | 22 | 22 |

Table 2: Results of the Rayleigh test for circular uniformly distributed data of the body orientation of the bees in all four environments and spatial categories. The statistical results of the z-value, p-value and significance level are given for each spatial area and the tested environmental condition.

| condition | area | z | p-value | significance level |
|---|---|---|---|---|
| Three objects | low clutter | 389.684 | 1.38e-170 | *** |
| | high clutter | 609.359 | 1.83e-270 | *** |
| | above clutter | 825.417 | 0 | *** |
| | below outside | 356.904 | 2.85e-16 | *** |
| | above outside | 137.680 | 1.03e-06 | *** |
| full-density | low clutter | 221.989 | 2.42e-97 | *** |
| | high clutter | 661.197 | 1.53e-29 | *** |
| | above clutter | 123.114 | 3.01e-54 | *** |
| | below outside | 548.840 | 1.16e-24 | *** |
| | above outside | 182.583 | 1.77e-82 | *** |
| half-density | low clutter | 770.239 | 0 | *** |
| | high clutter | 247.969 | 1.65e-11 | *** |
| | above clutter | 570.329 | 1.61e-249 | *** |
| | below outside | 212.517 | 0.119 | ns |
| | above outside | 900.812 | 0.0001 | ** |
| Outer ring | low clutter | 287.988 | 4.40e-126 | *** |
| | high clutter | 296.394 | 1.29e-13 | *** |
| | above clutter | 190.498 | 1.45e-83 | *** |
| | below outside | 188.734 | 6.26e-09 | *** |
| | above outside | 148.903 | 3.34e-07 | *** |

#### 2.1 Additional tables

The detailed statistical results of the Kuiper test and the v-test can be found in a separate file (BodyOrientation_Statistics.xlsx).
